# Mass-Sensitive Particle Tracking to Characterize Membrane-Associated Macromolecule Dynamics

**DOI:** 10.1101/2022.01.27.477846

**Authors:** Frederik Steiert, Tamara Heermann, Nikolas Hundt, Petra Schwille

## Abstract

Short-lived or transient interactions of macromolecules at and with lipid membranes, an interface where a multitude of essential biological reactions take place, are inherently difficult to assess with standard biophysical methods. The introduction of mass-sensitive particle tracking (MSPT) constitutes an important step towards a thorough quantitative characterization of such processes. Technically, this was made possible through the advent of interferometric scattering microscopy (iSCAT)-based mass photometry (MP). When the background removal strategy is optimized to reveal the two-dimensional motion of membrane-associated particles, this technique allows the real-time analysis of both diffusion and molecular mass of unlabeled macromolecules on biological membranes. Here, a detailed protocol to perform and analyze mass-sensitive particle tracking of membrane-associated systems is described. Measurements performed on a commercial mass photometer achieve time resolution in the millisecond regime and, depending on the MP system, a mass detection limit down to 50 kDa. To showcase the potential of MSPT for the in-depth analysis of membrane-catalyzed macromolecule dynamics in general, results obtained for exemplary protein systems such as the native membrane interactor annexin V are presented.

**SUMMARY:** This protocol describes an iSCAT-based image processing and single-particle tracking approach that enables the simultaneous investigation of the molecular mass and the diffusive behavior of macromolecules interacting with lipid membranes. Step-by-step instructions for sample preparation, mass-to-contrast conversion, movie acquisition, and post-processing are provided alongside directions to prevent potential pitfalls.

## INTRODUCTION

Once merely perceived as a barrier against the wide range of ambient physical conditions, biological membranes are nowadays considered as functional entities and catalytic platforms^1, 2^. Based on their ability to localize, amplify, and direct signals in response to membrane-associated macromolecule reactions, lipid interfaces constitute a crucial element for a wide variety of cellular processes such as membrane trafficking and signaling cascades^3–5^. Serving as nucleation event for the assembly of stable complexes, membrane attachment often relies on a dynamic equilibrium between membrane-associated and cytosolic forms of macromolecules and is hence of transient nature^6, 7^.

In spite of their great importance in biology, it has so far been challenging to develop methods that are able to provide access to the compositional, spatial, and temporal heterogeneities of membrane-associated macromolecule reactions in real-time^7, 8^. To resolve the underlying molecular processes, two experimental aspects are decisive: sufficient time resolution and single-particle sensitivity. Therefore, ensemble average techniques such as fluorescence recovery after photobleaching (FRAP) but also the much more sensitive fluorescence correlation spectroscopy (FCS) do have limitations, since they largely uncouple spatial from temporal information^9^. An important step towards the characterization of individual molecule dynamics has thus been the advent of single-particle tracking (SPT) in combination with highly sensitive microscopy. In particular, two SPT approaches have proven to be effective in this regard. Firstly, the utilization of fluorophores as labels and the corresponding fluorescence detection systems paved the way for nanometer precision and millisecond time resolution^10–12^. Secondly, scattering-based detection using gold nanoparticles improved both localization precision and time resolution to the subnanometer and microsecond range, respectively^13–16^. Despite the many advantages of both approaches and their significant contributions regarding the mechanistic understanding of mem- brane-associated systems^17, 18^, both techniques have so far been limited because they required labeling of the molecules of interest, potentially perturbing their native behavior and because they were insensitive to the molecular composition of membrane-associated particles^19, 20^.

Both these limitations have recently been overcome by the introduction of a novel interferometric scattering (iSCAT)-based approach termed mass photometry^21–23^. This technique allows the determination of in-solution mass distributions of biomolecules according to their interferometric scattering contrast when landing on a glass interface. However, for the detection and characterization of mobile molecules diffusing on lipid membranes, a more sophisticated image analysis approach had to be developed. This has meanwhile been successfully implemented and allows to detect, track and determine the molecular mass of single unlabeled biomolecules diffusing on a lipid interface^24, 25^. Referred to as dynamic mass photometry or mass-sensitive particle tracking, this technique now enables the assessment of complex macromolecule interactions by directly recording changes in the molecular mass of the tracked entities and hence opens up new possibilities for the mechanistic analysis of membrane-associated molecular dynamics.

Here, a detailed protocol for sample preparation, imaging, and the data analysis pipeline required for mass-sensitive particle tracking is presented. In particular, sample requirements and potential problems that may occur during measurement and analysis are discussed. Furthermore, the unparalleled potential to analyze membrane-interacting macromolecule systems is showcased through various representative results.

## PROTOCOL

### 1. MSPT sample preparation

1.1 Generation of multilamellar vesicles (MLVs)
  1.1.1 Calculate the quantity of chloroform-dissolved lipid(s) according to the desired lipid mixture and required suspension volume. NOTE: A final vesicle concentration of 4 mg/mL lipids is recommended for the resuspension (reaction) buffer.
  1.1.2 Pipette the calculated volume of lipids into a 1.5 mL glass vial using positive displacement pipettes equipped with glass tips.
  1.1.3 Evaporate the lipid solvent under a faint stream of nitrogen and constantly rotate the vial to ensure equal distribution of the lipids on the glass walls.
  1.1.4 Ensure full solvent evaporation by placing the vial below a steady stream of nitrogen for 15 min.
  1.1.5 Further remove residual traces of chloroform by vacuum drying in a vacuum desiccator for an additional hour.
  1.1.6 Rehydrate the lipid mixture in the desired reaction buffer and thoroughly vortex the suspension until the lipid film has been dissolved from the vial’s walls. NOTE: The reaction buffer should be chosen to ensure protein activity and stability. The reaction buffer used in this study contains 50 mM Tris-HCl (pH = 7.5), 150 mM KCl and 5 mM MgCl_2_. Note that any buffer used to dilute lipids or proteins needs to be filtered to remove interfering particulate impurities (see section 5).
1.2 Generation of small unilamellar vesicles (SUVs)
  1.2.1 For consecutive freeze-thaw cycles of the lipid resuspension (step 1.1.6), boil 500 mL of water in a beaker on a hot plate (between 70° C to 99° C) and prepare a container containing liquid nitrogen.
  1.2.2 Shock-freeze the lipid resuspension in the liquid nitrogen. Transfer the vial into the beaker with hot water until the solution is completely thawed. Repeat this freeze-thaw cycle 8-10 times or until the previously turbid mixture appears clear. CAUTION: Use proper safety garments and equipment such as goggles, gloves, and tweezers to prevent any direct contact with the liquid nitrogen, the frozen lipid vial, or the boiling water.
  1.2.3 For the generation of a monodisperse vesicle distribution, assemble a lipid-extruder and test its integrity with reaction buffer to ensure that it does not leak. NOTE: If leakage is observed, carefully re-assemble the lipid-extruder until no buffer spilling is evident.
  1.2.4 Extrude the lipid suspension for 37 passes through a nucleopore membrane with a pore size of 50 nm^26^. The number of passes should be uneven to ensure that the final SUV mixture crossed the nucleopore membrane and is hence free of lipid aggregates or multilamellar vesicles. The extruded vesicles will be used later to form supported lipid bilayers (see sections 6 and 7). NOTE: SUVs can likewise be formed by sonication of the rehydrated lipid mixture. However, preparation *via* extrusion provides a more monodisperse distribution of SUVs which facilitates vesicle rupture during supported lipid bilayer formation. Extruded vesicles can be stored in the fridge for a maximum of three days.

### 2. Cleaning of Microscope Slides

2.1 Distribute an equal number of microscope slides with thickness No. 1.5 (#1.5, 0.17 mm) and dimensions 24 mm x 60 mm, and 24 mm x 24 mm in polytetrafluorethylene (PTFE) microscope holders.
2.2 Transfer PTFE holders into beakers containing ultrapure water and sonicate them for 15 min at room temperature. NOTE: Depending on the beaker, water volume needs to be adjusted to completely cover the PTFE holder.
2.3 Use tweezers to remove the holders from the beaker and replace water with ultrapure isopropanol. Insert the holder into the beaker containing isopropanol and sonicate again for 15 min. NOTE: Depending on the beaker, the volume of the isopropanol needs to be adjusted to completely cover the PTFE holder.
2.4 Replace isopropanol with ultrapure water and sonicate the beaker containing the holders for 15 min.
2.5 Remove the PTFE holders from the beakers and blow-dry the microscope slides in the holder under a steady stream of nitrogen gas or compressed air. NOTE: Ensure proper cleaning of cover slides by using gloves, clean beakers, and sealing film to cover each beaker. Otherwise, residual dust might cause significant background fluctuations during MSPT measurements.

### 3. Hydrophilization of Microscope Slides

NOTE: To obtain a homogeneous and fluid-supported lipid bilayer, hydrophilization of slides is essential and must be carried out just before flow chamber assembly.

3.1 Place PTFE holders containing only #1.5, 24 mm x 60 mm microscope slides in a plasma cleaner with oxygen as a process gas and clean the microscope slides with plasma (parameters used in this work: 30% power, 0.3 mbar oxygen pressure for 30 s (see **Table of Materials** for details of plasma cleaner used).

NOTE: To obtain fluid membranes, plasma cleaning parameters such as power, oxygen pressure, and cleaning time must be adjusted for each instrument. For this purpose, the use of fluorescently labeled lipids is recommended to ensure membrane fluidity which can be quantified with fluorescence recovery after photobleaching (FRAP) experiments^27^. If the parameters are not optimized for the respective setup, membrane diffusion might be impaired due to reduced membrane fluidity.

### 4. Assembly of Flow Chambers

4.1 Before flow chamber assembly, keep the following components ready: cleaned microscope slides (#1.5, 24 mm x 24 mm), hydrophilized microscope slides (#1.5, 24 mm x 60 mm), aluminum foil, flat cardboard, scalpel, and double-sided tape.
4.2 Wrap the flat cardboard with aluminum foil.
4.3 Spread the cleaned #1.5, 24 mm x 24 mm microscope slides on the aluminum foil with sufficient distance between each other.
4.4 Attach double-sided tape strips to the upper and lower edges of the slides.
4.5 Excise each microscope slide with the scalpel, such that it can be removed from the aluminum foil. As a result, each slide should have stripes of double-sided tape attached to the upper and lower glass edge (see **Figure 1**).
4.6 Attach the #1.5, 24 mm x 24 mm slide with the two double-sided tape strips to the hydrophilized slide (#1.5, 24 mm x 60 mm) to form a flow path between the smaller and bigger microscope slide. NOTE: To ensure clean flow chambers, constantly wear gloves and ensure a dust-free workbench.

**Figure 1:**
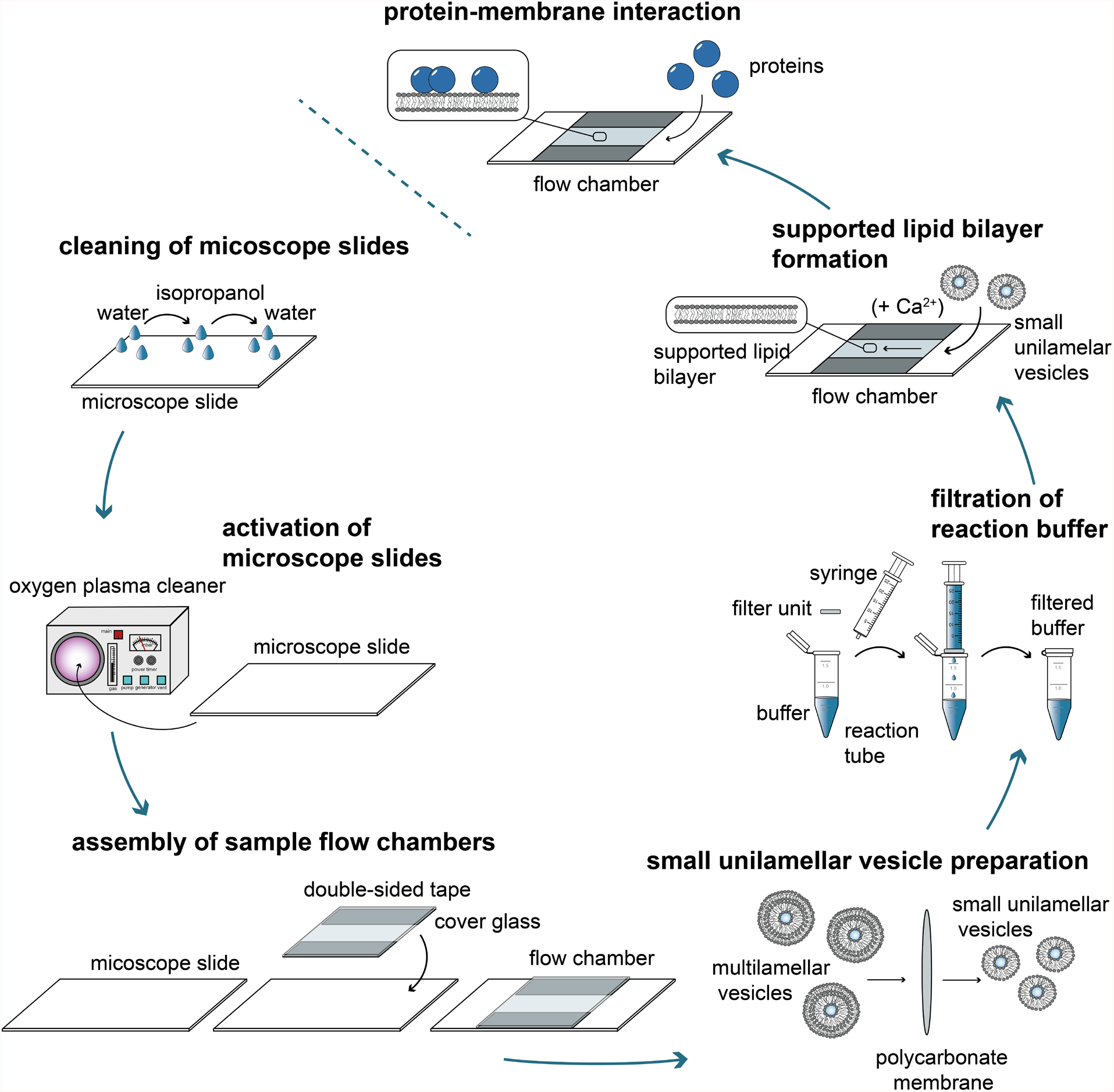
Process flow diagram of the individual steps required to analyze protein-membrane interactions with mass-sensitive particle tracking (MSPT). To prepare samples for MSPT meas- urements, glass cover slides must be thoroughly cleaned and activated with an oxygen plasma. After their assembly into sample flow chambers, small unilamellar vesicles (SUVs) are prepared for supported lipid bilayer (SLB) formation and all reaction buffers are filtered to reduce back- ground scattering. SUVs are added to form lipid bilayers in the flow chambers. Optionally, diva- lent cations such as Ca^2+^ ions may be added to the SUVs to promote vesicle rupture. At last, low concentrations of the protein of interest are flushed into the reaction chamber.

### 5. Filtration of Reaction Buffers

5.1 Sterile filter all reaction buffers through 0.45 μm cellulose acetate membranes to ensure minimal background signal during MSPT measurements.

NOTE: If the presence of nucleotides, such as ATP, is essential for successful experiment performance, be aware of a potential increase in background signal. It is recommended to only use minimal amounts that still ensure protein activity.

### 6. Supported lipid bilayer (SLB) formation

NOTE: It is recommended to perform the formation of supported lipid bilayers on the mass photometer to visually ensure successful vesicle spreading and the complete removal of unfused vesicles.

6.1 Dilute freshly extruded SUVs (see section 1 for more details) to a final concentration of 0.4 mg/mL in the required reaction buffer. Optionally, to promote vesicle rupture, add 2 mM CaCl_2_ to the vesicle suspension. NOTE: Divalent cations might cause the aggregation of some lipids such as PiP_2_. For mixtures containing such lipids refrain from using CaCl_2_ for the promotion of vesicle rupture or other divalent cations in the resuspension buffer. If required for the experiment, divalent cations can be added after the successful formation of the supported lipid bilayer.
6.2 Flush 50 μL of the vesicle suspension into the flow chamber (step 4) and incubate the chamber for 2 min. NOTE: Buffers, vesicles, or protein solutions can be flushed through the flow chamber with a small piece of soaking tissue. However, it is also possible to use a mechanical pump system.
6.3 Remove unfused vesicles through repeated (at least three times) washing of the flow chamber with each 200 μL of the reaction buffer. NOTE: Vesicles need to be thoroughly washed out of the flow chamber to ensure a stable background signal during MSPT measurements.

### 7. Generation of MSPT calibration curve

NOTE: To convert the contrast of detected particles into molecular mass, their signal needs to be calibrated using proteins of known sizes. It is recommended to adjust the standard protein size regime to cover the range of molecular masses expected for the system of interest.

7.1 Biotinylation of standard proteins with a cysteine residue
  7.1.1 Calculate the appropriate amount of maleimide-biotin for the standard protein according to the manufacturer’s instructions.
  7.1.2 Incubate the standard protein with the determined volume of maleimide-biotin for 1 h at room temperature.
  7.1.3 To remove unconjugated maleimide-biotin from the conjugated biotin-protein complex, perform size-exclusion chromatography on a column suitable for the protein of interest.
  7.1.4 Determine the protein concentration using a Bradford Assay. NOTE: To store the standard protein for further measurements, freeze the protein in single-use aliquots in liquid nitrogen and store them at -80 °C.
7.2 Measurement of standard proteins for the calibration curve
  7.2.1 In a flow chamber, form a supported lipid bilayer with 0.4 mg/mL extruded SUVs (see sections 1 and 6 for more details) containing 0.01 mol% (v/v) Biotinyl Cap PE (1,2-dioleoyl-sn-glycero- 3-phosphoethanolamine-N-cap biotinyl).
  7.2.2 Add 50 μL of 2.5 nM divalent streptavidin to the bilayer in the flow chamber and incubate for 10 min. NOTE: Divalent streptavidin has been expressed and purified as outlined in Howarth et al.^28^. Tetravalent streptavidin can likewise be used. However, the use of divalent streptavidin can reduce the possible reaction stoichiometries between biotinylated lipids and standard proteins that are conjugated to a biotin moiety in order to facilitate the assignment of species.
  7.2.3 Remove unbound divalent streptavidin with 100 μL of reaction buffer.
  7.2.4 Add 50 μL of 100 nM biotin-conjugated standard protein to the bilayer in the flow chamber and incubate for 2 min. NOTE: Depending on the biotinylation efficiency and whether di- or tetravalent streptavidin is used, the optimum concentrations of biotin-conjugated standard protein and streptavidin may vary.
  7.2.5 Perform MSPT measurement according to the details outlined in section 8. CAUTION: Imaging conditions need to be identical for both sample and calibration standards.

### 8. Imaging

8.1 SLB formation and sample preparation
  8.1.1 As described in more detail in section 6, introduce SUVs of the desired lipid mixture (25 μL) to the sample flow chamber and form a supported lipid bilayer. Thoroughly wash the chamber with reaction buffer (three times 100 μL) to remove all unfused vesicles.
  8.1.2 Add 50 μL of the protein of interest to the sample chamber. NOTE: As MSPT is a single-particle method, protein concentration has to be kept in the pM to nM range to allow for undisturbed particle detection and tracking.
8.2 Video acquisition
  8.2.1 Set the desired imaging conditions such as the size of the field of view (FOV), frame rate, exposure time, and acquisition time in the acquisition software. NOTE: The following settings have proven to work for MSPT on a commercial mass photometer (**Table of Materials**): FOV of 128 pixels x 35 pixels, a frame rate of 1 kHz, resulting in roughly 200 frames per second after subsequent 5-fold frame averaging, and an exposure time of 0.95 ms.
  8.2.2 Adjust the focus automatically or manually. If necessary, move the FOV to a position with a homogenous membrane using the lateral control.
  8.2.3 Create a project folder and start recording the movie. Upon completion of the recording, specify a file name in the dialog prompted by the acquisition software. The movie is then automatically saved to the project folder as a MP file for subsequent analysis. NOTE: Record at least three replicates in different flow chambers to ensure the integrity of individual membranes and reproducibility of results. The movie duration can be set in advance and depends on the type of experiment. In most cases, an acquisition time between 5 and 7 min is recommended. CAUTION: By default, movie recordings are compressed before being saved to reduce storage space. However, file compression needs to be turned off to enable custom data analysis as described in this protocol. Details on how to turn off file compression can be found in the manufacturer’s user manual.

### 9. Data analysis

NOTE: The data analysis pipeline is accompanied by an interactive Jupyter notebook. The Jupyter notebook and associated custom-written Python modules necessary to perform the MSPT analysis outlined below are available in a public repository: https://github.com/MSPT-toolkit/MSPT-toolkit. For detailed instructions on the analysis below, readers are referred to ***MSPT analysis*.*ipynb*** accessed using the above link.

9.1 Video processing
  9.1.1 Remove dominant static scattering of light with the pixel-wise background estimation algorithm (using ***image_processing*.*mp_reader*** function). To apply the background removal, choose the option ***continuous_median*** for the parameter ***mode*** and set an appropriate length for the sliding median window (***window_length***) in notebook section B.1. Optionally, save the movies after background removal to be used for particle detection and trajectory linking (***save_processed_movies***). NOTE: Adjust the window size (***window_length***) to values between 101 and 2001 depending on the particle density on the membrane, the expected diffusion coefficient, acquisition frame rate, and the required processing speed. CAUTION: The background removal strategy works well if the membrane is not too densely packed and if the diffusion of the particles is sufficiently fast (i.e., each pixel is most of the time not occupied with a particle). Otherwise, the contrast of the particles will be systematically underestimated as they cannot be properly distinguished from the background signal. This can be compensated for by increasing the median window size at the cost of computational speed. However, be aware that setting the window size too large may negatively influence the output due to sample drift. A visual inspection of the processed videos is crucial.
  9.1.2 Detect particles and their respective position throughout the movie using the function ***particle_fitting*.*particle_fitter_parallel*** (notebook section B.2). Tune the sensitivity of particle detection with the threshold parameter (***thresh***, notebook section B.1), which is used to highlight candidate spots by image binarization. The effect of varying threshold parameters on spot detection sensitivity can be examined in a separate notebook (***Movie visualization*.*ipynb***). The results of particle detection are automatically saved to CSV files in a subdirectory of the movie file. NOTE: Setting the threshold parameter arbitrarily low (for example, for movies taken with the used mass photometer), a threshold parameter below 0.0055 is not recommended as candidate spots will be dominated by spurious noise and hence prolong processing time.
  9.2 Link particles in consecutive frames into trajectories using the Python package *trackpy* (v.0.5.0)^29^. The trajectory linking is performed on the fly after spot detection. As a result, an additional CSV file containing the trajectory information is stored in a subdirectory of the particle detection CSV file. Remove trajectories with too few points using the parameter ***minimum_trajectory_length*** (notebook section B.1) to enable a robust determination of diffusion coefficients. For detailed explanations regarding the other parameters of *trackpy* functions, please refer to *trackpy*’s documentation.
9.3 Trajectory analysis
  9.3.1 In notebook section C.1, specify the frame rate (***frame_rate***) and pixel size in nm (***pixel_size***), which were used for movie acquisition. Create a list of CSV files containing the trajectory information returned by *trackpy* (see step 9.2) with the ***trajectory_analysis*.*get_csv_files*** function (notebook section C.2). Additionally, specify an output filename for the HDF5 container, which is used to store the fitting results on disk (notebook section C.3). Analyze all trajectories with the ***trajectory_analysis*.*fit_trajectories*** function in notebook section C.4 which iterates through the list of CSV files. This function uses jump distance distribution (JDD)^30^ and mean squared displacement (MSD)^31^ analysis to estimate the diffusion coefficient of each trajectory.
  9.3.2 Convert the median contrast of each trajectory to its corresponding mass using the contrast-mass relationship obtained from the MSPT calibration (see section 7). Specify the slope (***slope***) and y-intercept (***offset***) of the calibration line, which relates iSCAT contrast to molecular mass (function ***trajectory_analysis*.*apply_calibration***; see notebook section C.5). This function adds a column with the trajectory’s median mass to each data frame in the HDF5 container.
  9.3.3 Evaluate the apparent particle density on the membrane with the function ***trajectory_analysis*.*membrane_density***, which returns the median density value in terms of detected particles and present trajectories during each frame (see notebook section C.6) as additional columns in the data frame. NOTE: As a portion of particles will be lost during both the detection and trajectory linking process, the actual particle densities may be higher. For reliable results regarding particle densities as well as mass histograms, visually inspect representative movie snapshots to verify that the measurement conditions are plausible for single-particle tracking (see section 9.1.1).

### 10. Data visualization

10.1 Illustrate the correlation of mass and diffusion coefficient with two-dimensional kernel density estimation (KDE), which is based on the Python package *fastkde* (v.1.0.19; https://pypi.org/project/fastkde/). To generate the plot, specify the HDF5 file containing the MSPT results (see section 9.3.1 and notebook section D.1) and select a single (D.2) or a concatenated data frame (D.3) as input data for the ***plotting*.*generate_2D_KDE*** function (notebook section D.4). NOTE: Each plotted dataset should contain ideally more than 1000 trajectories for a reliable 2DKDE.

## RESULTS

Following the detailed protocol herein for the preparation of supported lipid bilayers (SLBs) in flow chambers (**Figure 1**), one can clearly recognize a speckle-like pattern in the native view of all displayed conditions (**Figure 2**). This effect is caused by the surface roughness of the glass, which generally dominates the scattering signal and leads to visually indistinguishable conditions (glass, glass with SLB, or glass with SLB and attached proteins). The presence of vesicles, however, is clearly distinct due to the vesicles’ large scattering cross-section and enables the observation of vesicle rupture and fusion into homogeneous membranes (**Figure 2b** and **Supplementary Movie 1**). When removing the static scattering signal of the glass surface with a ratiometric approach that emphasizes the dynamic elements within the field of view^24,25^, one can uncover unlabeled proteins diffusing on the membrane (**Figure 2d**) while an empty SLB (**Figure 2c**) or glass itself (**Figure 2a**) appear as a noisy image.

**Figure 2:**
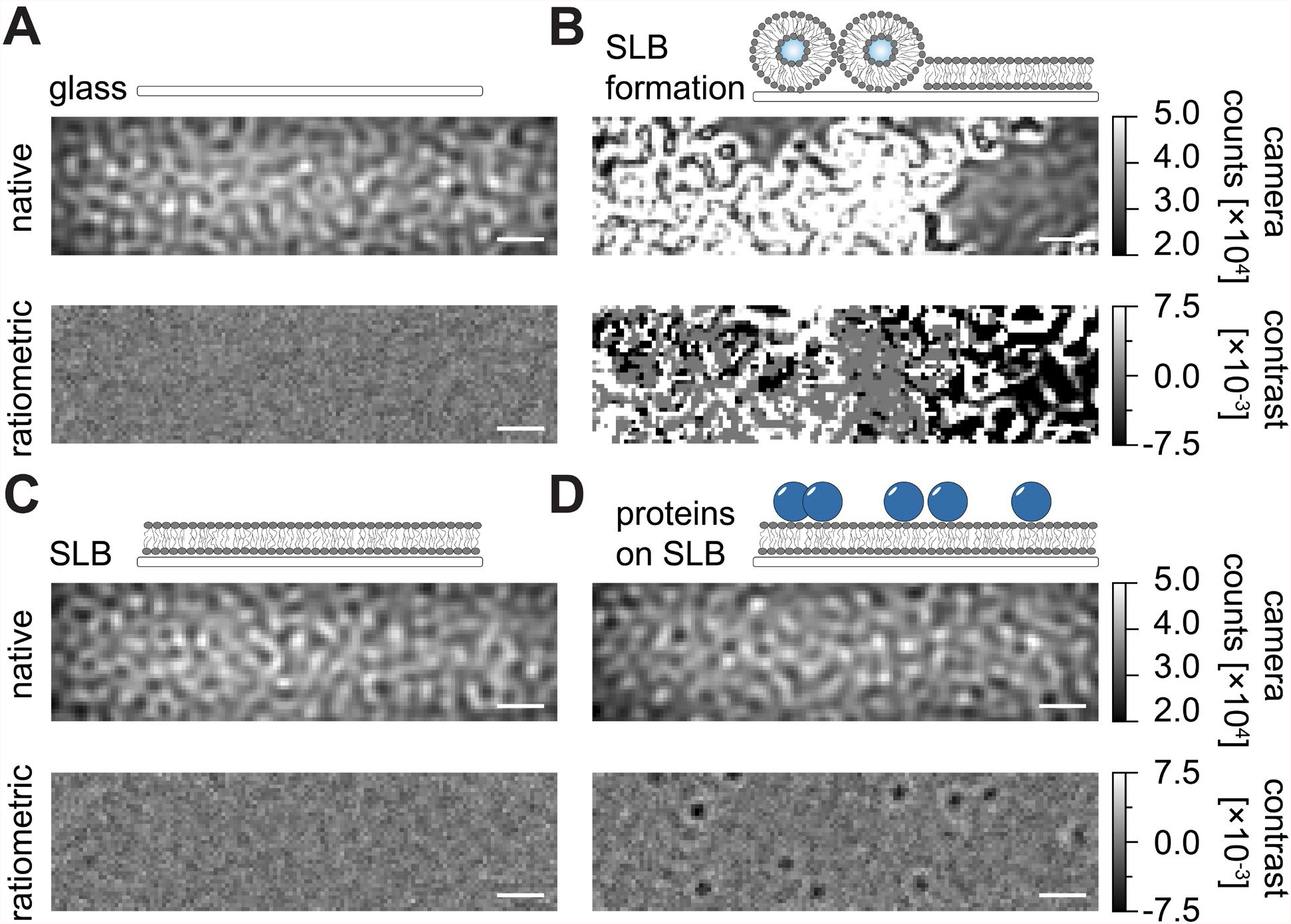
Native and ratiometric view of exemplary surfaces relevant for MSPT measurements. Representative images of the surface roughness of a glass cover slide (**a**), during the formation of a supported lipid bilayer (**b**), with an intact supported lipid bilayer (**c**) and of exemplary pro- teins reconstituted on an SLB (**d**). All four examples are displayed in the native mode, which can be accessed during the measurement itself, and as processed ratiometric images after median- based background removal. Scale bars represent 1 μm. For data analysis (see accompanied Jupy- ter notebook, protocol section 9), the following parameters were used: median window size (*win- dow_length*) = 1001.

The inherent background of MSPT measurements can be locally estimated by dividing each pixel value through the median of *n* preceding and succeeding pixels of the movie at the same image position (**Figure 3**). As a result, macromolecules appear as isotropic point-spread functions (PSFs) whose motion on the membrane can be observed, tracked, and quantified. In fact, the availability of both contrast and dynamic behavior enables the direct relation of a particle’s molecular size to its respective diffusive behavior, all without the need for labeling the particle. Nevertheless, to interpret the iSCAT contrast determined during MSPT experiments, it is essential to perform a calibration that translates the signal amplitude into molecular mass. This can be achieved by attaching biomolecules of known mass to an SLB *via* a biotin-streptavidin-biotin complex (**Figure 4a**). As an exemplary strategy, one can use biotinylated variants of bovine serum albumin (BSA), protein A (prA), alkaline phosphatase (AP), and fibronectin (FN), which bind to streptavidin (STP) that itself is bound to biotin-containing lipids (Biotinyl Cap PE) in the membrane. As displayed in **Figure 4a**, the increasingly pronounced contrast of these exemplary macromolecules reflects the increasing molecular weight of the respective biotinylated standards. By assigning each peak of the contrast histograms (**Figure 4b**) to the corresponding mass of the standard protein’s oligomer state, a linear relationship between contrast and mass is revealed^21,22^ and can subsequently be used for the analysis of unknown macromolecule systems (**Figure 4c**).

**Figure 3:**
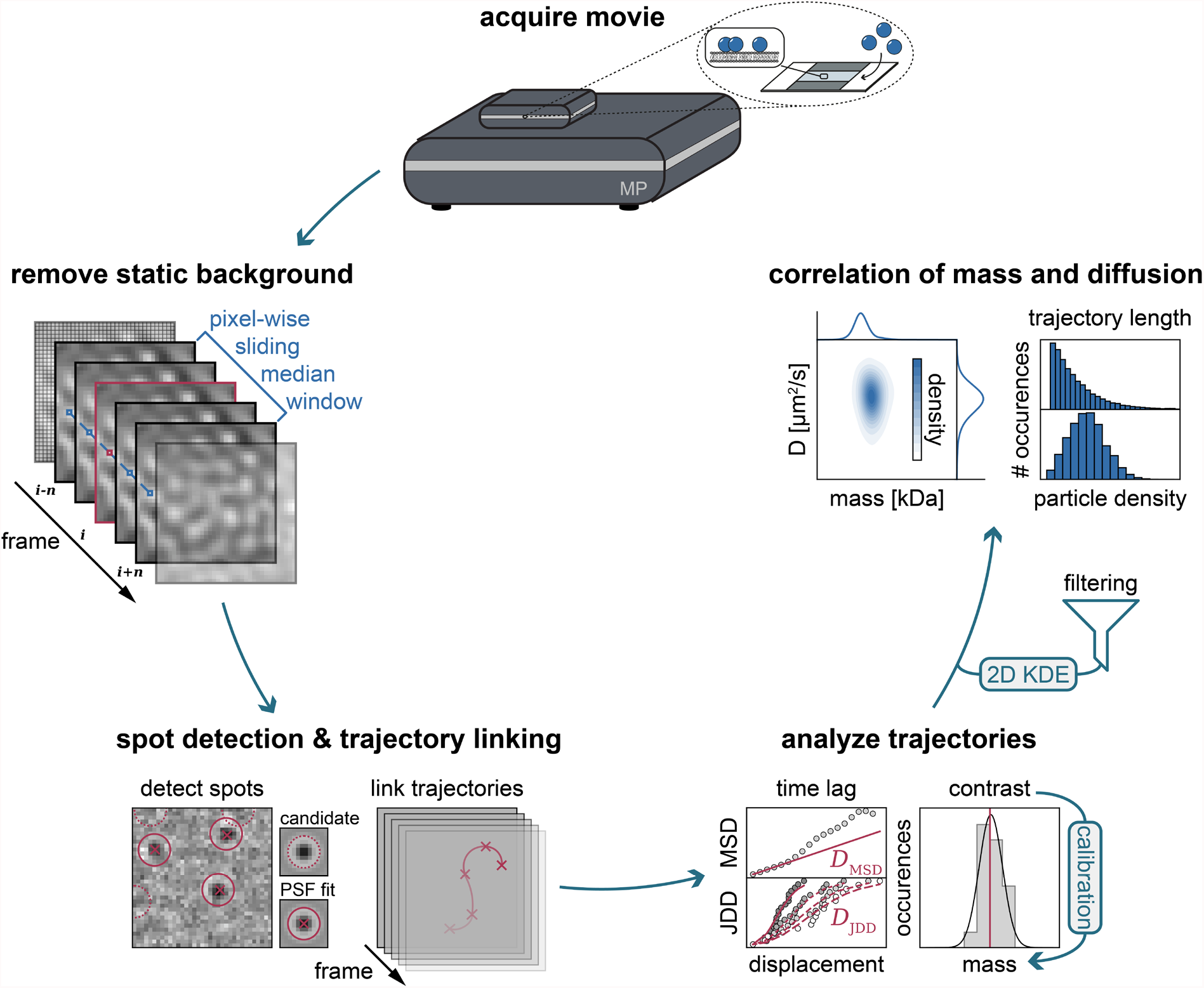
Step-by-step diagram of the stages required for MSPT data collection and analysis. After data acquisition for the sample of interest on the mass photometer, movies are processed to remove the static background through a pixel-wise sliding median approach. Thereafter, can- didate particles are identified and fitted by a point spread function (PSF) prior to their linking into particle trajectories. To enable the determination of the diffusion coefficient for each particle, mean squared displacement (MSD) or jump distance distribution (JDD) analysis is employed. At this stage, contrast values can be transformed into molecular masses according to the contrast- mass-relation determined through the calibration strategy. As a final step, trajectories can be filtered based on their length or membrane particle density and visualized by two-dimensional kernel density estimation (2D-KDE).

**Figure 4:**
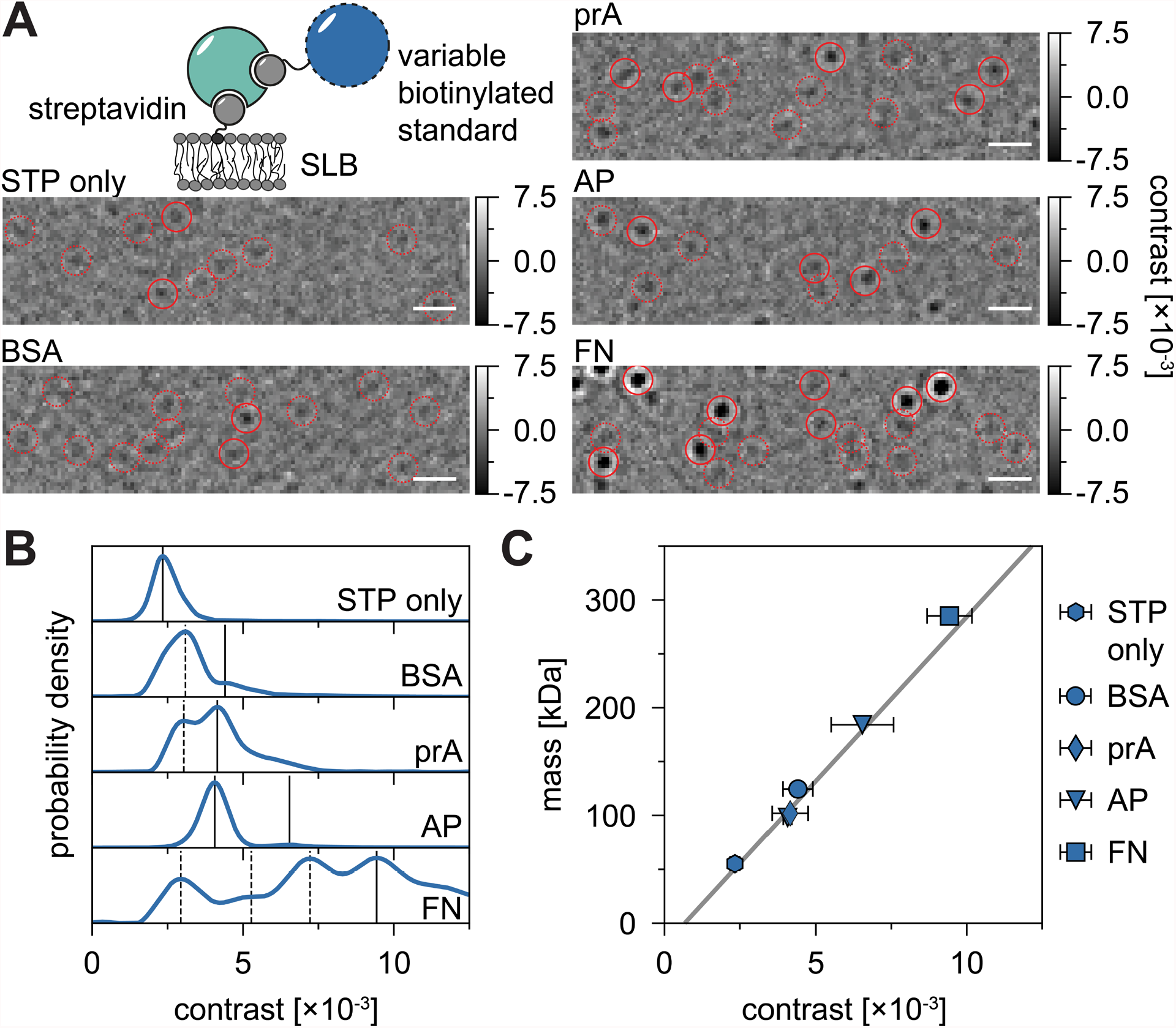
Calibration of the mass-to-contrast relation for MSPT measurements. **(a)** Representa- tive ratiometric frames obtained for exemplary streptavidin-standard protein complexes diffus- ing on a supported lipid bilayer containing a small percentage of biotinylated lipids (70:29.99:0.01 mol% - DOPC:DOPG:Biotinyl Cap PE). As model molecular weight standards, monovalent strep- tavidin^28^ (STP only) or divalent streptavidin^28^ in complex with either biotinylated bovine serum albumin (BSA), biotinylated protein A (prA), biotinylated alkaline phosphatase (AP), or biotinyl- ated fibronectin (FN) are shown. Candidate spots (dotted circle) and successful particle detec- tions (solid circle) are highlighted in red. The scale bars represent 1 μm. **(b)** Probability density distributions of contrast values obtained for the five model standard proteins. All displayed data represent pooled distributions of three independent experiments per condition: STP only n = 82,719; BSA n = 9,034; prA n = 22,204; AP n = 69,065 and FN n = 71,759 trajectories. As compared to particle numbers determined for membranes with proteins, the number of particles detected on an empty bilayer is negligible at moderate membrane densities (**Supplementary Figure 1**). Contrast peaks considered for mass calibration are marked through continuous lines while dashed ones represent oligomer states not considered. **(c)** Contrast-to-mass calibration curve, derived from peak contrasts in panel (**b**) and the respective sequence masses of the complexes. Error bars display the standard error of the peak locations estimated by bootstrapping (100 resamples with 1000 trajectories each). For data analysis (see accompanied Jupyter notebook, protocol section 9), the following parameters were used: median window size (*window_length*) = 1001 frames, detection threshold (*thresh*) = 0.00055, search range (*dmax*) = 4 pixels, memory (*max_frames_to_vanish*) = 0 frames, minimum trajectory length (*minimum_trajectory_length*) = 7 frames (STP only), 9 frames (BSA/FN), 15 frames (prA), 10 frames (AP).

A good example demonstrating the applicability and capabilities of MSPT to analyze molecular weights and hence study oligomer states and oligomerization events is the consideration of biotinylated aldolase and biotinylated IgG (**Figure 5**). Aldolase is commonly reported to be a homotetramer^32^. However, the mass distribution resolved by MSPT features four distinct peaks, which highlights the presence of multiple populations (**Figure 5a**). While the first minor peak corresponds to unoccupied streptavidin and can be expected due to the configuration in this kind of experiment, aldolase complexes with only two subunits (2SU) or 6 subunits (6SU) can likewise be detected (**Figure 5b**). Interestingly, tetra- and hexameric aldolase-streptavidin complexes display a reduced diffusion coefficient when compared to dimeric aldolase and streptavidin alone, indicating an increased viscous drag, e.g., *via* the attachment of a second biotinylated lipid to the streptavidin. Similarly, biotinylated IgG exhibits three peaks in the mass distribution, with the first peak again matching the mass of a single streptavidin. The mass of the most abundant peak corresponds to the mass of one light and one heavy chain (1SU), i.e., one-half of an IgG antibody.

**Figure 5:**
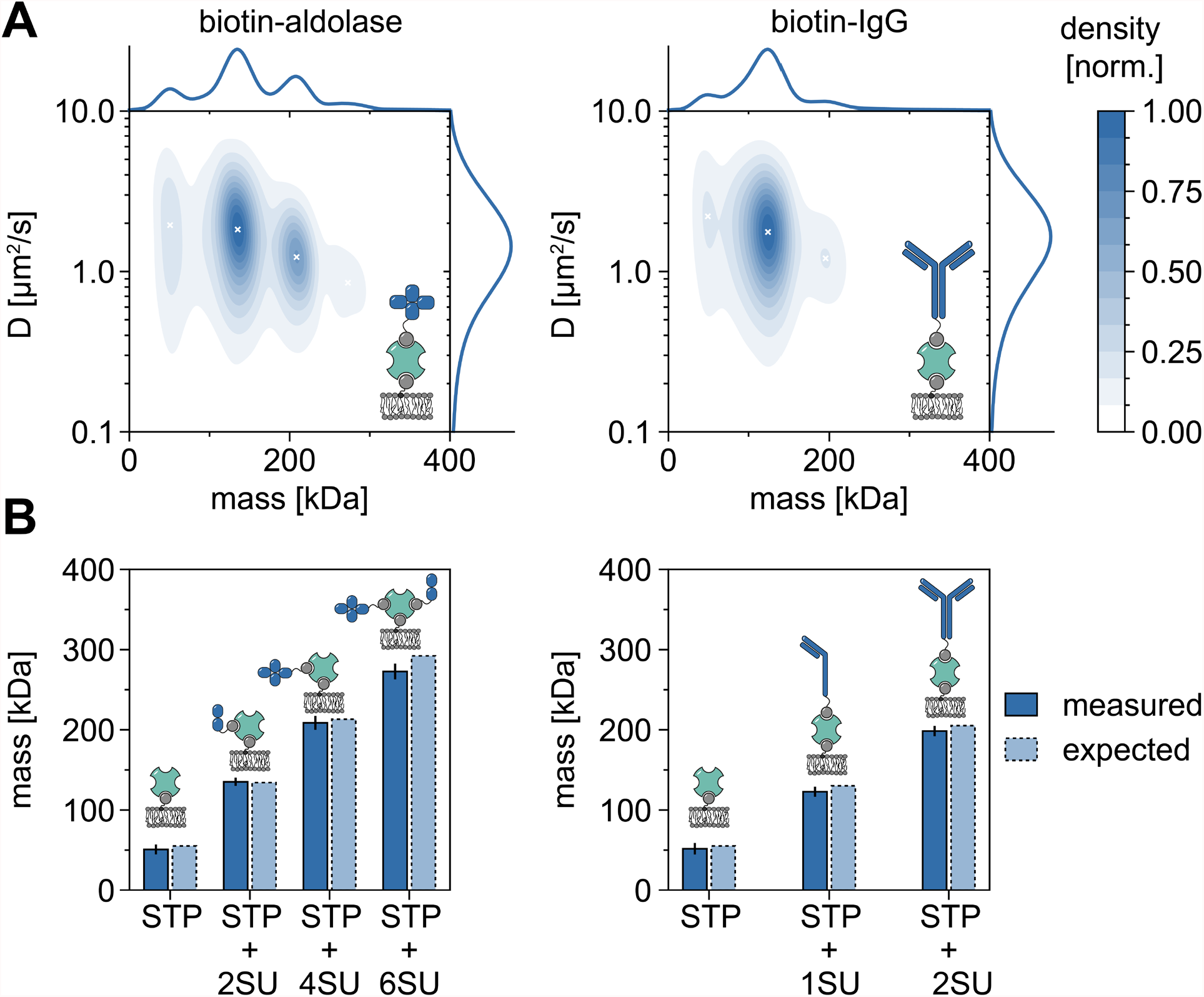
Deciphering oligomer states of membrane-associated proteins. **(a)** 2D kernel density estimations of both mass and diffusion coefficient of tetravalent streptavidin in complex with biotinylated aldolase (left panel) or with a biotin-modified goat anti-Rabbit IgG antibody (right panel). Reconstitution of both complexes has been performed on a supported lipid bilayer con- taining DOPC, DOPG, and Biotinyl Cap PE in a ratio of 70:29.99:0.01 mol%, respectively. In total, 116,787 trajectories of three independent replicates have been included for the streptavidin- aldolase complex (particle density of 0.1 μm^-2^) and 348,405 for the streptavidin-IgG composite (particle density of 0.1 μm^-2^). Only particles with a track length of at least 5 frames were included. Marginal probability distributions of both molecular mass (top) and diffusion coefficient (right) are presented. The black x in both panels mark the respective local maxima of the KDE. **(b)** Com- parison of determined oligomer masses for the complex of tetravalent streptavidin with biotin- modified aldolase (left panel) or biotinylated IgG (right panel) with, according to the sequence masses, expected molecular weights. The abbreviation SU is introduced on behalf of the protein of interests’ subunit. Error bars display the standard error of the peak locations estimated by bootstrapping (100 resamples with 1000 trajectories each). For data analysis (see accompanied Jupyter notebook, protocol section 9), the following parameters were used: median window size (*window_length*) = 1001 frames, detection threshold (*thresh*) = 0.00055, search range (*dmax*) = 4 pixels, memory (*max_frames_to_vanish*) = 0 frames, minimum trajectory length (*mini- mum_trajectory_length*) = 5.

The full antibody with two identical halves (2SU) is detected in about 11% of the cases. The decrease of the diffusion coefficient with increasing complex sizes indicates interactions of the streptavidin with more than one biotinylated lipid or additional drag caused by the attached IgG, or both.

Besides the sole analysis of membrane-dependent oligomer states, MSPT also confers the particular advantage of correlating the diffusive behavior of a macromolecule of interest with its oligomer state. Representative results for this type of analysis are shown for annexin V (AnV) and cholera toxin subunit B (CTxB) which bind to dioleoylphosphatidylserine (DOPS) or glycosphingolipids (GM1), respectively, incorporated in the membrane (**Figure 6a**). Both kernel density estimations (KDEs) feature unimodal distributions of mass and diffusion, indicating a single abundant species with similar diffusive behavior. The peak position of molecular mass and diffusion coefficient was found to be 49.8 ± 2.2 kDa and 1.4 ± 0.1 μm^2^/s, respectively, for AnV as well as 62.7 ± 3.1 kDa and 0.4 ± 0.1 μm^2^/s, respectively, for CTxB. The measured diffusion coefficients are comparable to previously reported values obtained from high-speed AFM and FRAP^33, 34^. The slightly reduced mass as compared to the mass of the expected macromolecule (52 kDa for an AnV trimer, 65 kDa for a CTxB pentamer) may indicate the presence of smaller complexes with fewer subunits in the ensemble. While the mass difference between the proteins is small and close to the specified detection limit of the microscope (≈50 kDa), their diffusion coefficients differ considerably. In an equimolar mixture, for instance, by comparing the diffusion of the mixture to the distribution of AnV and CTxB alone, one can conclude that AnV is more abundant on the membrane than CTxB (**Figure 6b**). However, if the concentration of CTxB is doubled as compared to the concentration of AnV, the equilibrium is shifted towards CTxB as the predominant protein on the membrane. As illustrated for mixtures of AnV and CTxB, MSPT not only allows to discriminate membrane-associated macromolecules according to their molecular weight but also enables the discrimination of different macromolecule populations according to their diffusive behavior.

**Figure 6:**
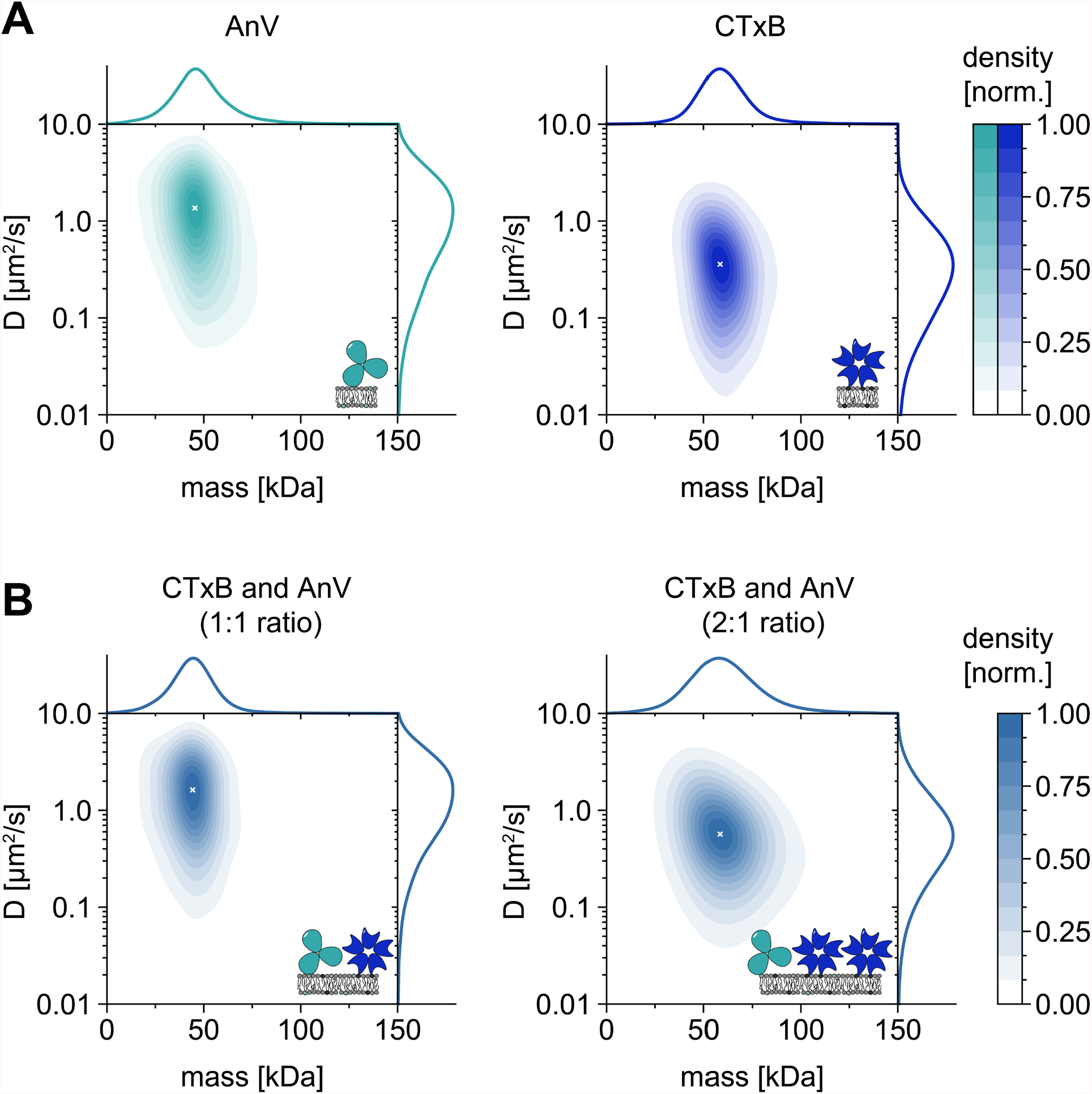
Dissolving the diffusive behavior of the native membrane-interacting proteins annexin V (AnV) and cholera toxin subunit B (CTxB). **(a)** 2D kernel density estimations of both mass and diffusion coefficient of annexin V (left panel) and cholera toxin subunit B (right panel). For AnV and CTxB membrane reconstitution, lipid compositions of 80:20 mol% DOPC to DOPS and 99.99:0.01 mol% DOPC to GM1 have been used, respectively. In total 206,819 trajectories of three independent replicates have been included for AnV (particle density of 0.1 μm^-2^) and 142,895 for CTxB (particle density of 0.2 μm^-2^). **(b)** 2D kernel density estimations of CTxB and AnV mixtures in a ratio of 1:1 (left panel) or 2:1, respectively. Reconstitution of protein mixtures was performed on a supported lipid bilayer containing DOPC, DOPS and GM1 lipids in a ratio of 80:19.99:0.01 mol%. In total 42,696 trajectories of three independent replicates have been included for the 1:1 mixture (particle density of 0.1 μm^-2^) and 264,561 trajectories for the 2:1 ratio (particle density of 0.3 μm^-2^). For both (**a**) and (**b**) only particles with a track length of at least 5 frames were included. Marginal probability distributions of both molecular mass (top) and diffusion coefficient (right) are presented. The white x in each panel marks the respective global maximum of the KDE. For data analysis (see accompanied Jupyter notebook, protocol section 9), the following parameters were used: median window size (*window_length*) = 1001 frames, detection threshold (*thresh*) = 0.00055, search range (*dmax*) = 4 pixels, memory (*max_frames_to_vanish*) = 0 frames, minimum trajectory length (*minimum_trajectory_length*) = 5.

As with all microscopy techniques, some experimental requirements are crucial in order to achieve the desired quality of data. An important example in this context is thoroughly cleaned coverslips. In general, this is considered a prerequisite for microscopy-related single-molecule experiments, but MSPT is particularly sensitive to sample impurities. The increased scattering originating from the glass surface of uncleaned coverslips prevents any quantitative iSCAT measurement. Notably, even residual dirt or dust particles on insufficiently cleaned glass can cause notable image distortions, recognizable as bright spots in the native imaging mode (**Figure 7a**). Although these defects are removed by the background estimation due to their static nature, accurate determination of a particle’s contrast might be impaired and thus negatively influences its quantitative analysis. Another common problem encountered in MSPT experiments is remaining vesicles that either float (encircled in orange) through the field of view or unfused vesicles that are stuck (encircled in blue) at a specific position on the membrane and appear as large pulsating scatterers **(Figure 7b**). To minimize their occurrence and interference with movie acquisition, it is recommended to thoroughly wash the SLB before adding the protein and to use freshly prepared mixtures of small unilamellar vesicles (SUVs) and divalent cations.

**Figure 7:**
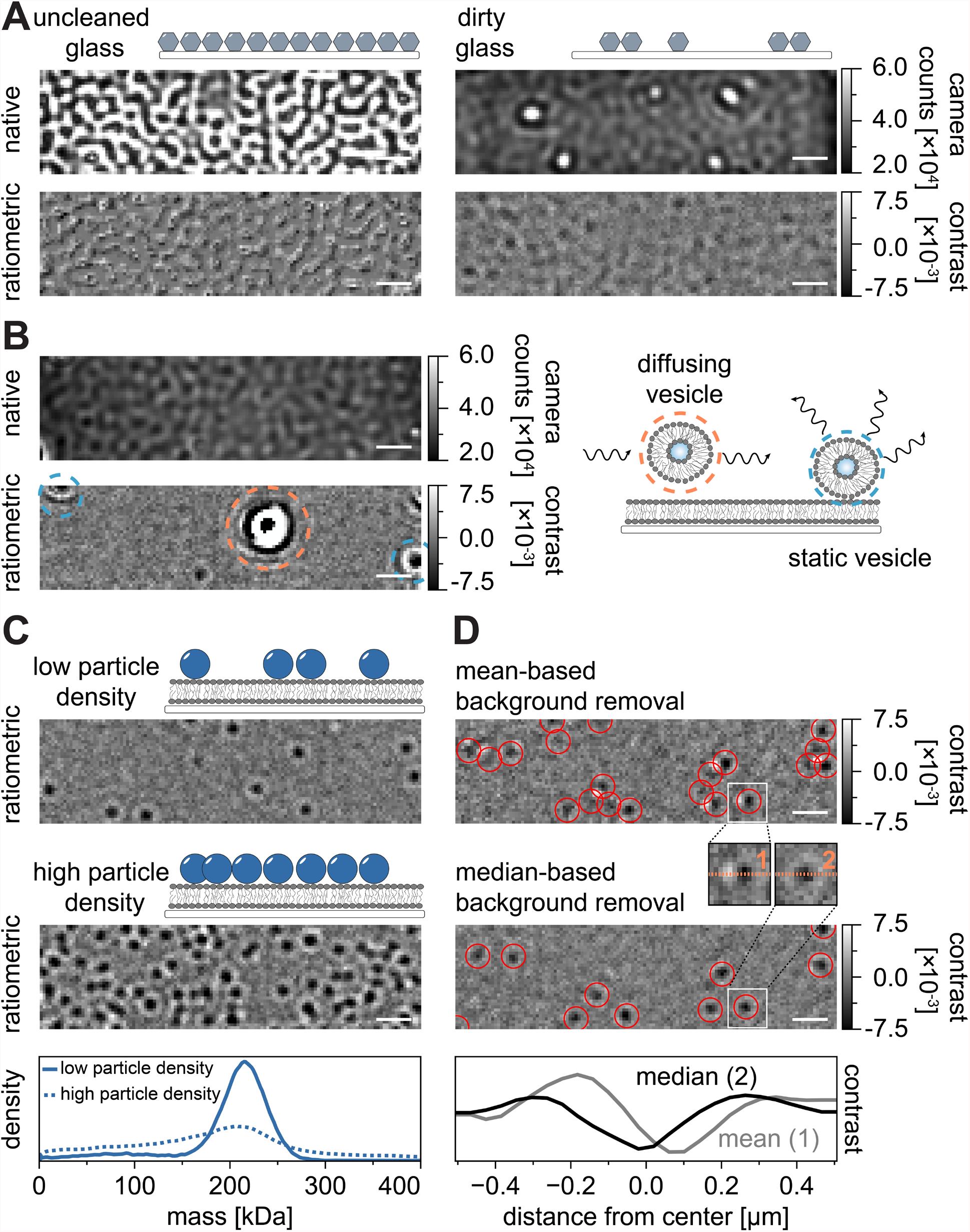
Potential complications in the course of MSPT measurements or during data analysis. **(a)** Representative images of the surface roughness displayed in both the native and the processed (median-based background removal) ratiometric view of an uncleaned cover glass slide. In both cases, bright spots constitute residual surface impurities that impede artifact-free measurements. **(b)** Exemplary images of residual vesicles in the field of view after insufficient membrane washing. Both static (highlighted in blue) and diffusing (highlighted in orange) vesicles will impair measurement quality either due to pulsating and wiggling or due to their directional movement, respectively. **(c)** As a single-particle technique, MSPT requires low particle densities (representative image, upper panel) to enable proper linking and mass-determination of each particle. In the case of high membrane-particle densities (middle panel), particle fitting is impaired which affects mass-determination (see lower panel). **(d)** Representative ratiometric images of particles diffusing on a membrane interface after either mean-based (upper panel) or median-based background removal. For diffusing particles, the mean-based background removal strategy produces distorted images of the particle’s PSF as can be seen in the small insets between the upper and the middle panel. In contrast, undistorted particle PSFs can be obtained through the median-based approach. Lower panel: Comparison of line profiles through the center of the PSF obtained after mean- or median-based background removal. For all native and ratiometric images displayed in this figure, the scale bars represent 1 μm. For data analysis (see accompanied Jupyter notebook, protocol section 9), the following parameters were used: median window size (*window_length*) = 1001 frames, detection threshold (*thresh*) = 0.00055, search range (*dmax*) = 4 pixels, memory (*max_frames_to_vanish*) = 0 frames, minimum trajectory length (*minimum_trajectory_length*) = 5.

One factor that must be taken into account for the design of mass-sensitive particle tracking experiments is the density of macromolecules associated with the membrane interface. High particle densities on the membrane can in fact cause problems for two reasons: i) The linking of particle detections from consecutive frames into trajectories becomes ambiguous and hence increases the likelihood of errors and misjudged diffusion coefficients. ii) The mass of particles, which is extracted from the amplitude of their corresponding PSF fit, becomes systematically underestimated and mass peaks broaden because it becomes increasingly difficult to separate the static background signal from the dynamic particle signal (**Figure 7c**). Currently, visual assessment of data quality in the process of acquiring MSPT videos is difficult on the available commercial microscopes because the implemented ratiometric view in the acquisition software is using the background removal established for mass photometry^21^ instead of the median-based algorithm described here and in references ^24, 25^ (**Figure 7d**). The mean-based continuous background removal used to visualize landing molecules in mass photometry causes diffusing particles to appear as dark fronts with bright tails which makes the spots appear highly anisotropic and interferes with PSF fitting during the detection procedure. Thus, the use of the implemented meanbased image processing in the acquisition software is unsuitable for the analysis of diffusing biomolecules on membranes.

## DISCUSSION

The presented protocol extends mass photometry^21^, a technique that analyzes the mass of single biomolecules adsorbing on glass, to an even more versatile tool capable of simultaneously measuring the mass and diffusion of unlabeled membrane-interacting biomolecules. This analysis extension is achieved through the implementation of a modified background removal strategy adapted to the lateral motion of molecules^24, 25^. In general, the background removal is of utmost importance for iSCAT-based approaches, since the strong scattering of the glass surface roughness represents the main analysis impediment, and the accurate determination of each pixel’s local background is essential for quantification of particle mass and location. Besides the imageanalysis adapted to particle motion, subsequent particle detection, trajectory linking, and data analysis complete the novel expansion of MP into mass-sensitive particle tracking (MSPT).

In general, thoroughly cleaned glass cover slides and a clean working environment are critical requirements for the successful performance of MSPT experiments. Due to the absence of macromolecule labeling, the acquired signal is inherently nonselective. Clean samples, as well as proper sample handling, are hence crucial to ensure that observations cannot be misinterpreted. In particular, when molecules of low molecular weight are examined, control measurements of protein-free membranes are recommended to assess background contributions (**Supplementary Figure 1**). Besides the inclusion of control measurements, it is thus recommended to follow the preparation steps displayed in **Figure 2** for each flow chamber. When combined, these safety measures will ensure that the detected signal originates from the biomolecule of interest and not, for example, a contaminated flow chamber, buffer, or membrane.

Besides the precautions regarding the experimental design, care needs to also be taken during MSPT image processing. During video processing, the value for three parameters should be chosen carefully to ensure correct results: i) the length of the median window for background removal, ii) the threshold for particle detection, and iii) the maximum search radius during the linking assignment. A larger median window (i) generally facilitates the separation of diffusing particles from the superimposed quasi-constant background. However, for too large window sizes sample drift will eventually become noticeable and diminish the accuracy of the background estimation. Optimal settings heavily depend on sample properties and measurement conditions. Nevertheless, a value of 1001 can be used as a robust starting point. The threshold parameter (ii) must be tuned depending on the lowest molecular mass expected in the sample. A value below 0.005 is not recommended for measurements taken with the mass photometer used in this study. To speed up analysis times, higher values can be chosen if a sample with high molecular weight is expected. The search radius in trajectory linking (iii) specifies the maximum radial distance in pixels in which the particle’s shifted location will be searched for in consecutive frames. It should be adapted to the fastest particle in the sample, and, if favored, an adaptive search range (see documentation of *trackpy*) could be used instead to reduce computation time. Especially during the initial phase of a project re-analyzing the movies with varying parameters is recommended to validate the obtained results.

In light of the single-molecule nature of MSPT, it should be avoided to measure at high membrane particle densities as those can interfere with accurate contrast and mass determination. It has been shown that densities below one particle per squared micrometer are favorable for MSPT measurements^24^. An additional consideration is the expected diffusion coefficients in the sample. Although applicable to a broad range of diffusion coefficients, MSPT has a lower limit of accessible diffusion coefficients. Local confinement to a region of few pixels during a significant portion of the median window period merges the particle with the static background. For the imaging conditions used in this protocol, measurement of diffusion coefficients below 0.01 μm^2^/s is not recommended. At this diffusion speed, for example, the mean squared displacement of a particle during the median window half-size is about 4 pixels and hence of similar size as the extent of the PSF. As a consequence, the static background estimate is likely to contain signal contributions from the particle itself, which results in an apparently reduced contrast of the particle until it eventually approaches the noise level. However, macromolecule diffusion coefficients ranging between 0.05 and 10 μm^2^/s can clearly be resolved.

To further extend the range of MSPT applications, one can envision an advancement of the median-based background algorithm through the elimination of pixels that are temporarily occupied with a particle, or by sample drift correction enabling larger median window sizes. Both approaches would alleviate the problems regarding measurements at high particle densities and slow diffusion. Improvements in terms of lower mass sensitivity are on the horizon with a new generation of mass photometers, which may provide access to biomolecules smaller than 50 kDa. Therefore, future MSPT experiments will be able to study single-molecule dynamics and mem- brane-related interactions for an even wider range of membrane mimicries such as cushioned bilayers and macromolecular systems.

## ACKNOWLEDGMENTS

We sincerely appreciate the support from Philipp Kukura, Gavin Young, and the Refeyn software team and acknowledge their assistance by sharing parts of the image analysis code. We thank the Cryo-EM MPIB Core Facility for providing access to the commercial Refeyn mass photometer. F.S. gratefully acknowledges the support and funding granted by Jürgen Plitzko and Wolfgang Baumeister. T.H. and P.S. received funding through the Deutsche Forschungsgemeinschaft (DFG, German Research Foundation) - Project-ID 201269156 – SFB 1032 (A09). N.H. was supported by a DFG return grant HU 2462/3-1. P.S. acknowledges support through the research network MaxSynBio via the joint funding initiative of the German Federal Ministry of Education and Research (BMBF) and the Max Planck Society.

## DISCLOSURES

The authors have nothing to disclose.

## Notes

### Competing Interest Statement

The authors have declared no competing interest.

