## Supplementary Information for "Mass-Sensitive Particle Tracking to Characterize Membrane-Associated Macromolecule Dynamics"

**TITLE**

**AUTHORS AND AFFILIATIONS**

Frederik Steiert[1,2], Tamara Heermann[1], Nikolas Hundt[3,*] and Petra Schwille[1,*]

[1]Department of Cellular and Molecular Biophysics, Max Planck Institute of Biochemistry, Am Klopferspitz 18, 82152 Planegg, Germany
[2]Department of Physics, Technical University Munich, 85748 Garching, Germany.
[3]Department of Cellular Physiology, Biomedical Center (BMC), Ludwig-Maximilians-Universität München, Großhaderner Str. 9, 82152 Planegg, Germany

* Corresponding Authors:
Nikolas Hundt
Petra Schwille

Email Addresses of co-authors:
Frederik Steiert
Tamara Heermann
Nikolas Hundt
Petra Schwille

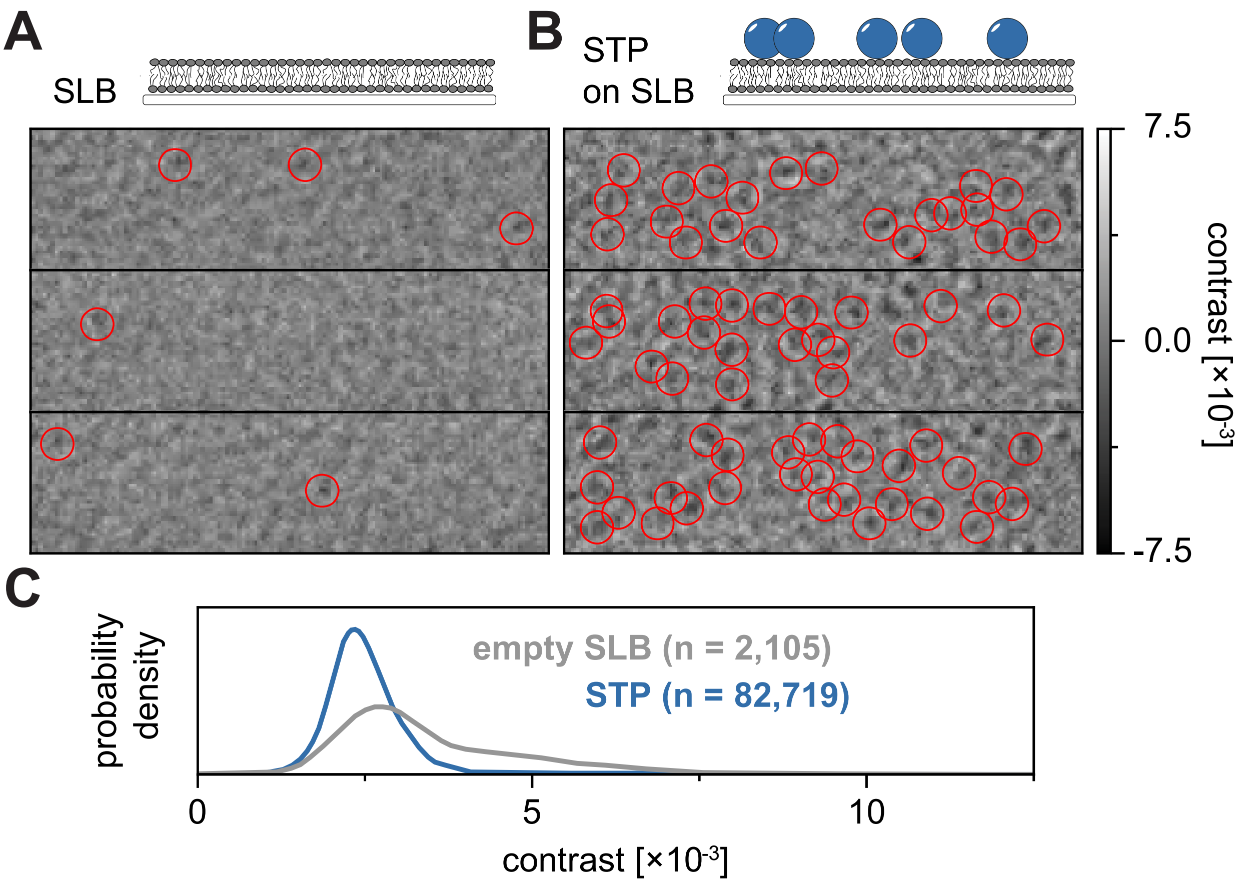


**Supplementary Figure 1: Comparison of protein-free and occupied membranes.** Representative images of an intact supported lipid bilayer before (**a**) and after (**b**) the addition of purified streptavidin (STP). Candidate spots that were successfully fit to the model PSF are encircled in red. (**c**) Contrast probability distributions of particles detected on an empty membrane (membrane background, grey) and on a bilayer with diffusing streptavidin particles (blue). Both probability distributions represent the pooled data of three independent experiments with identical movie acquisition and analysis parameters. For data analysis (see accompanied Jupyter notebook, protocol section 9), the following parameters were used: median window size (*window_length*) = 1001 frames, detection threshold (*thresh*) = 0.00055, search range (*dmax*) = 4 pixels, memory (*max_frames_to_vanish*) = 0 frames, minimum trajectory length (*minimum_trajectory_length*) = 7 frames.

**Supplementary Movie 1**: Exemplary movie showing the rupture and fusion of vesicles into a homogeneous membrane recorded with the mass photometer. Image processing median window size (*window_length*) = 1001 frames. Scale bar: 1 µm. Camera counts range: black = 16892; white = 65408.

**Supplementary Movie 2**: Exemplary movies showing the diffusion of annexin V (top) and biotinylated aldolase (bottom) complexes on a bilayer as obtained from MSPT measurements. Image processing median window size (*window_length*) = 1001 frames. Scale bar: 1 µm. Interferometric scattering contrast range: black = -0.0075; white = 0.0075.
